# Is the Success of Adaptive Therapy in Metastatic Castrate-Resistant Prostate Cancer Influenced by Cell-Type-Dependent Production of Prostate-Specific Antigen?

**DOI:** 10.1101/2021.10.21.465292

**Authors:** Monica Salvioli, Len Vandelaer, Katharina Schneider, Rachel Cavill, Kateřina Staňková

**Affiliations:** Department of Engineering Systems and Services, Delft University of Technology, Delft, The Netherlands; Department of Data Science and Knowledge Engineering, Maastricht University, Maastricht, The Netherlands

**Keywords:** Metastatic castrate-resistant prostate cancer, prostate-specific antigen, adaptive therapy, tumor composition, game theory, qualitative resistance

## Abstract

Prostate-specific antigen (PSA) is the most common serum marker for prostate cancer. It is used to detect prostate cancer, to assess responses to treatment and recently even to determine when to switch treatment on and off in adaptive therapy protocols. However, the correlation between PSA and tumor volume is poorly understood. There is empirical evidence that some cancer cell types produce more PSA than others. Still, recent mathematical cancer models assume either that all cell types contribute equally to PSA levels, or that only specific subpopulations produce PSA at a fixed rate.

Here, we compare time to competitive release of the PSA-based adaptive therapy protocol by Zhang et al. with that of the standard of care based on continuous maximum tolerable dose under different assumptions on PSA production. In particular, we assume that androgen dependent, androgen producing, and androgen independent cells may contribute to the PSA production to different extents.

Our results show that, regardless the assumption on how much each type contributes to PSA production, the time to competitive release is always longer under adaptive therapy than under the standard of care. However, in some cases, e.g., if the androgen-independent cells are the only PSA producers, adaptive therapy protocol by Zhang et al. cannot be applied, because the PSA value never reaches half of its initial size and therefore therapy is never discontinued.

Furthermore, we observe that in the adaptive therapy protocol, the number of treatment cycles and their length strongly depend on the assumptions about the PSA contribution of the three types. Our results support the belief that a better understanding of patient-specific PSA dynamics will lead to more successful adaptive therapies.

## 1. Introduction

Prostate-specific antigen (PSA) is an enzyme produced by prostate epithelial cells, both normal and cancerous ones [2]. The PSA level in blood is influenced by many factors, including the age of the patient, the ethnic group, the size of prostate, the presence of prostate cancer and its stage and tumor volume [22, 24, 23]. For this reason, the precise correlation between the PSA level and the tumor volume remains poorly understood [33, 25, 3]. Nevertheless, PSA is currently the most widely used serum marker to diagnose, stage and monitor prostate cancer and to assess responses to treatment [2, 12, 29, 4]. With the advent of new treatment strategies such as adaptive therapy (AT), which modulates the treatment depending on the response of the specific patient [14, 42], getting more information on tumor response to treatment and progression became even more crucial.

In a recent clinical trial applying AT in patients with metastatic Castrate-Resistant Prostate Cancer, decision-making was entirely PSA-based: The patients were given maximum tolerable dose of abiraterone until their PSA level dropped to 50% or less of its initial value. At this point, abiraterone was discontinued until PSA returned to the starting value. Each patient received this formula of cycling on and off in response to their PSA levels until radiographic progression [42]. While the cycles on and off abiraterone varied widely from patient to patient, this trial demonstrated that the adaptive dosing extends the time to progression more than twice, compared to the standard of care applying abiraterone continuously at the maximum tolerable dose (MTD) (median time to progression of about 30 months compared to about 14 months for standard of care) [1, 41].

The AT protocol was determined through a game-theoretical model of metastatic Castrate-Resistant Prostate Cancer, where three competing cancer cell types were identified: *T*^+^ cells requiring exogenous androgen to survive, *T*^*P*^ cells producing testosterone due to the upregulation of the enzyme CYP17A and *T*^−^ cells which are androgen-independent [39, 40]. Zhang et al. (2017) assumed that each of these cell types produces one unit of PSA and that 50% of the PSA decays out of the blood serum per unit time:

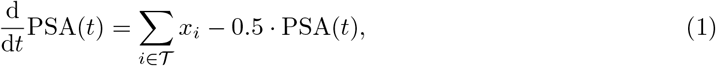

with 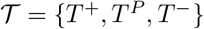 and *x*_*i*_ being the number of cells of the corresponding type [42].

The PSA dynamics have been explored in detail in many mathematical models. For instance, West et al. (2019) used the same assumptions of Zhang et al. (2017) and extended the formula to model four different cancer cell types [37]. Hansen et al. (2020) kept the 50% decay rate but assumed that each cell type produces two units of PSA per time unit [19]. As it is unclear how precisely the PSA level decays, Cunningham et al. (2018) and Cunningham et al. (2020) did not assume any decay rate but assumed that PSA simply measures 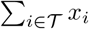 [10, 11].

Hirata et al. (2010) considered three slightly different cell types: androgen-dependent cells, androgen-independent cells resulting from reversible changes, and androgen-independent cells arising from irreversible changes of genetic mutations. Still they assumed that each type produces one unit of PSA without any decay, similarly to other works [20, 21, 30, 34, 32, 17, 35].

Brady-Nicholls et al. (2020 and 2021) proposed a model of prostate cancer stem cells and nonstem prostate cancer cell dynamics to simulate the observed PSA response patterns [5, 6]. In this approach, only differentiated non-stem cancer cells are simulated to produce PSA with a fixed rate, while stem cells do not. The model has been calibrated and validated to patient-specific data of two different clinical trials (Bruchovsky et al., 1990; Zhang et al., 2017) with PSA decay while on treatment and PSA increase during treatment holidays [7, 42].

In vitro experiments by Gustavsson et al. (2005), who cultured an androgen-dependent human prostate cancer cell line until the appearance of an androgen-independent sub-line and measured the corresponding PSA secretion, suggest that cell types may contribute to PSA production differently [18]. This would mean that the PSA dynamics, as introduced in [42, 10], might not reflect the actual tumor burden and that a more precise estimation of the PSA could be derived by accounting for the heterogeneity of the tumor cell population.

Consistent with this finding, we build on the model by Zhang et al. (2017), Cunningham et al. (2018), and Cunningham et al (2020), and assume that the three cell types can produce different amounts of PSA and explore different scenarios. In particular, we are interested in addressing and modelling the consequences of this assumption on the effectiveness of AT.

In the next section, we introduce the model and the parameters. Following [39, 42, 10, 11], we consider three categories of patients, based on their response to the treatment: best responders, responders and non-responders. In Section 3, we measure the superiority of AT over continuous MTD for each category, assuming that the different cell types contribute to PSA production to different extents. Section 4 concludes this paper by summarizing its outcomes and discussing limitations and future research.

## 2. Model

We use the Lotka-Volterra competition model by [42, 10, 11] to describe the interactions between the testosterone-dependent *T*^+^, the testosterone-producer *T*^*P*^ and the testosterone-independent *T*^−^ cell types under abiraterone therapy. The instantaneous rate of change in the population size of each cell type 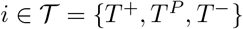 is:

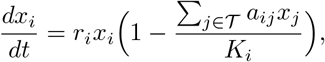

where *r*_*i*_ represents the growth rates, *K*_*i*_ the carrying capacities and *a*_*ij*_ the coefficients of the competition matrix

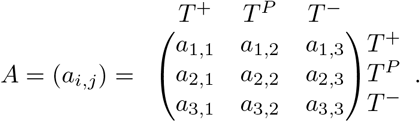

As in Zhang et al. (2017), Cunningham et al. (2018), and Cunningham et al. (2021), we set the growth rates to 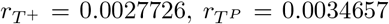 and 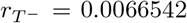, which are derived from the measured doubling times of representative cell lines [42, 10, 11, 9].

Following [42, 10, 11], we assume that abiraterone reduces the ability of *T*^+^ and *T*^*P*^ cells to acquire testosterone and we model this effect as a reduction in the carrying capacity of these cell types. In particular, abiraterone diminishes the ability of *T*^*P*^ cells to exploit the CYP17A pathway to convert cholesterol into androgens and therefore inhibits the production of testosterone. For this reason, in the absence of treatment the carrying capacity of the *T*^*P*^ cells is set to 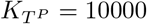, while under treatment it is reduced to 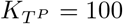. As the *T*^+^ cells rely on the endogenous testosterone produced by the *T*^*P*^ cells, following [42, 10, 11] we assume that their carrying capacity is a linear function of the density of the 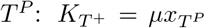, where *μ* = 1.5 in the absence of therapy and *μ* = 0.5 under therapy. As the *T*^−^ cells are not affected by abiraterone, their carrying capacity is always 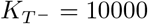.

Each competition coefficient *a*_*i,j*_ describes the effect of cells of type *j* on the growth rate of cells of type *i*. The intra-cell type coefficients are set to *α*_*i,i*_ = 1. Zhang et al. (2017), You et al. (2017), Cunningham et al. (2018), and Cunningham et al. (2020) assumed that the inter-cell type coefficients have values from the set {0.4, 0.5, 0.6, 0.7, 0.8, 0.9}. They distinguished 22 cases, which they group into three categories, depending on the frequency of *T*^−^ cells at the equilibrium [42, 39, 10, 11]:

- **Best responders**: twelve cases with a competition matrix promoting the absence of *T*^−^ and high frequencies of both *T*^+^ and *T*^*P*^. Like Cunningham et al. (2018) we use the following representative competition matrix for this category to explore model predictions [10]:

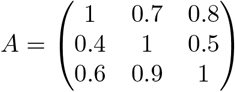
- **Responders**: four cases with competition matrices resulting in low frequencies of *T*^−^ at initiation of therapy. Following [10] for this category we use this representative competition matrix:

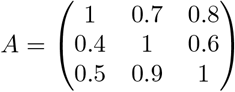
- **Non-responders**: six cases with a competition matrix resulting in high equilibrium frequencies of *T*^−^ (≥ 20%). As in [10], for this category we use the following representative competition matrix:

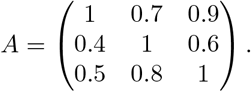

The initial cell counts for each category are taken from [10] and reported in Table 1.

**Table 1:** Initial cell counts, taken from [10].

| | $x_{T^+}(0)$ | $x_{T^P}(0)$ | $x_{T^-}(0)$ |
| --- | --- | --- | --- |
| Best responder | 606.06 | 757.58 | $1.94 \cdot 10^{-10}$ |
| Responder | 560.36 | 747.59 | 47.10 |
| Non-responder | 319.63 | 707.76 | 273.97 |

As opposed to [42, 10, 11], where the PSA level at a certain time *t* is assumed to correspond to the total number of cancer cells at that time up to some decay, here we assume that the three cell types can produce different amounts of PSA, so that the PSA level at a certain time *t* corresponds to:

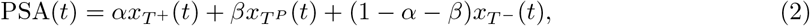

with 0 ≤ *α* ≤ 1 and 0 ≤ *β* ≤ 1 − *α*. For each representative case, we compare the outcome under continuous MTD to the outcome under AT, where the treatment is administered until the PSA drops to half of its initial value, then discontinued and readministered only when the PSA recovers to its initial level. Following common [11, 38], we refer to adaptive therapy only if the treatment is discontinued at least once.

In our case studies, we measure the success of the treatment through the time to competitive release (TCR), defined as the time at which *T*^−^ cells become the majority of the tumor composition. Following [42], we define this time as

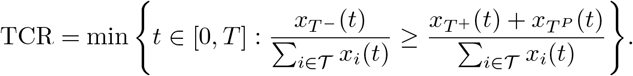

In our model, the treatment is applied even after reaching the TCR.

## 3. Results

In the following sections, we compare the effectiveness of the standard of care applying MTD with AT under different assumptions on the PSA production. We present results for the three patient categories: best responder, responder, and non-responder. In most scenarios, we consider four different assumptions on the PSA production: 1) all cell types contribute equally to PSA production, i.e., 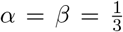, 2) only *T*^+^ cells produce PSA, i.e., *α* = 1, *β* = 0, 3) only *T*^*P*^ cells produce PSA, i.e., *α* = 0, *β* = 1, and 4) only *T*^−^ cells produce PSA, i.e., *α* = 0, *β* = 0. However, in other scenarios we also consider all possible values of *α* and *β.*

### 3.1. Best responders

Figure 1 illustrates the population size of the three different cell types *T*^+^, *T*^*P*^, and *T*^−^, as well as the total cell count when applying MTD (Figures 1A and 1E) or AT (Figures 1B-1D). Here, we focus on the best responder scenario. The TCR is highlighted with a yellow dot and the yellow-shaded area covers the time after competitive release, when the treatment protocol is continued but strategically the treatment has already failed.

**Figure 1:**
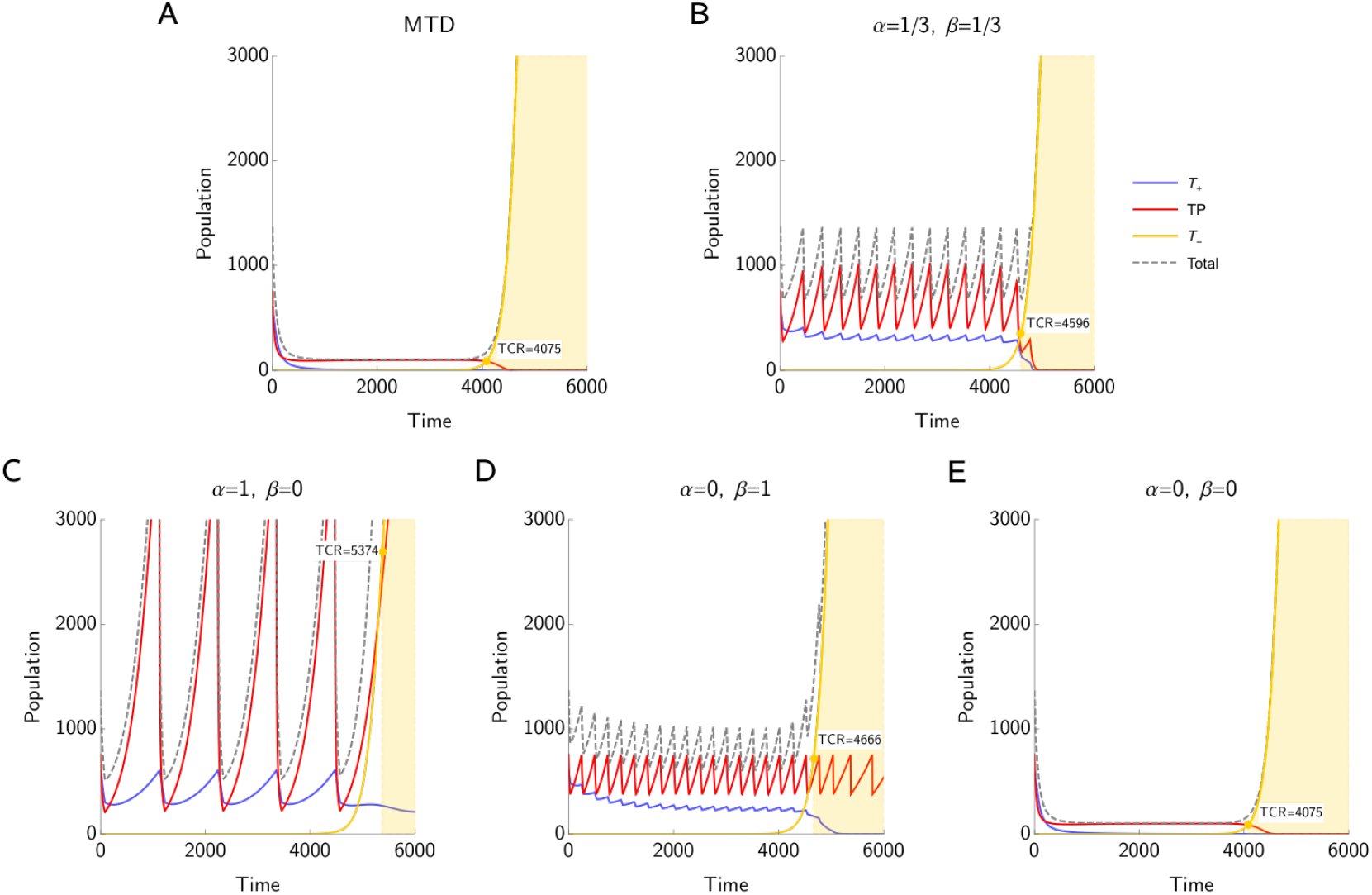
Best responders: Time to competitive release under MTD (A and E) and AT for different values of *α* and *β* (B-D). For all the values of *α* and *β* considered here, AT is better than the standard of care based on MTD. If *α* = *β* = 0, i.e., *T*^−^ cells are the only PSA producers, we cannot apply AT because the PSA never drops to half of its initial value before TCR. The number of treatment cycles in the AT protocol, as well as their length, vary depending on *α* and *β*. It is important to note that treatment is continued after reaching TCR. After TCR, we observe oscillations in the population sizes of *T*^*P*^ only if *T*^*P*^ supports PSA production, i.e., if *β >* 0.

We observe that in all cases AT can increase TCR compared to applying MTD. However, if *α* = *β* = 0, we cannot apply AT, as the *T*^−^ cells are not targeted by the treatment and, thus, the PSA level never drops to half of its initial value. Table 2 demonstrates the superiority of AT over MTD: Under AT the TCR is increased by 32%, 14%, and 13%, if only *T*^+^ cells are contributing to the PSA production, only *T*^*P*^ cells are contributing to PSA production, and all three types are contributing to the PSA production equally, respectively.

**Table 2:** Best responders: TCR under MTD and under AT including the TCR improvement percentage depending on different assumptions on PSA production. Applying AT increases TCR in all cases. We observe the highest improvement in the TCR for *α* = 1, *β* = 0.

| Parameter values | TCR under MTD | TCR under AT | Improvement |
| --- | --- | --- | --- |
| $\alpha = 1; \beta = 0$ | 4075 | 5374 | 32% |
| $\alpha = 0; \beta = 1$ | 4075 | 4665 | 14% |
| $\alpha = \frac{1}{3}; \beta = \frac{1}{3}$ | 4075 | 4596 | 13% |
| $\alpha = 0; \beta = 0$ | 4075 | N/A | N/A |

The number of treatment cycles as well as their length vary depending on *α* and *β*. In particular, when the PSA production is supported mainly by the *T*^*P*^ cells, we observe shorter treatment cycles leading to higher frequency in the oscillations of the cancer population size. This is caused by the fact that *T*^*P*^ cells are directly targeted by the treatment, resulting in an immediate response in the PSA level if their contribution to the PSA production is high. *T*^+^ cells are only influenced by the treatment via the *T*^*P*^ cells and thus, there is a small delay in the drop of the PSA level, which in turn leads to longer treatment cycles.

If *α* = 0 and *β* = 1, we observe strong oscillations in the population size of *T*^*P*^ cells. As long as enough *T*^*P*^ cells are present and their contribution to the PSA production is high enough, the PSA level can be influenced by the treatment and thus, the AT protocol will lead to continuing treatment cycles after TCR.

While in Figure 1 we focus on the population size dynamics for a few selected values of *α* and *β*, Figure 2 shows a heat map of TCR for all possible values of *α* and *β*.

**Figure 2:**
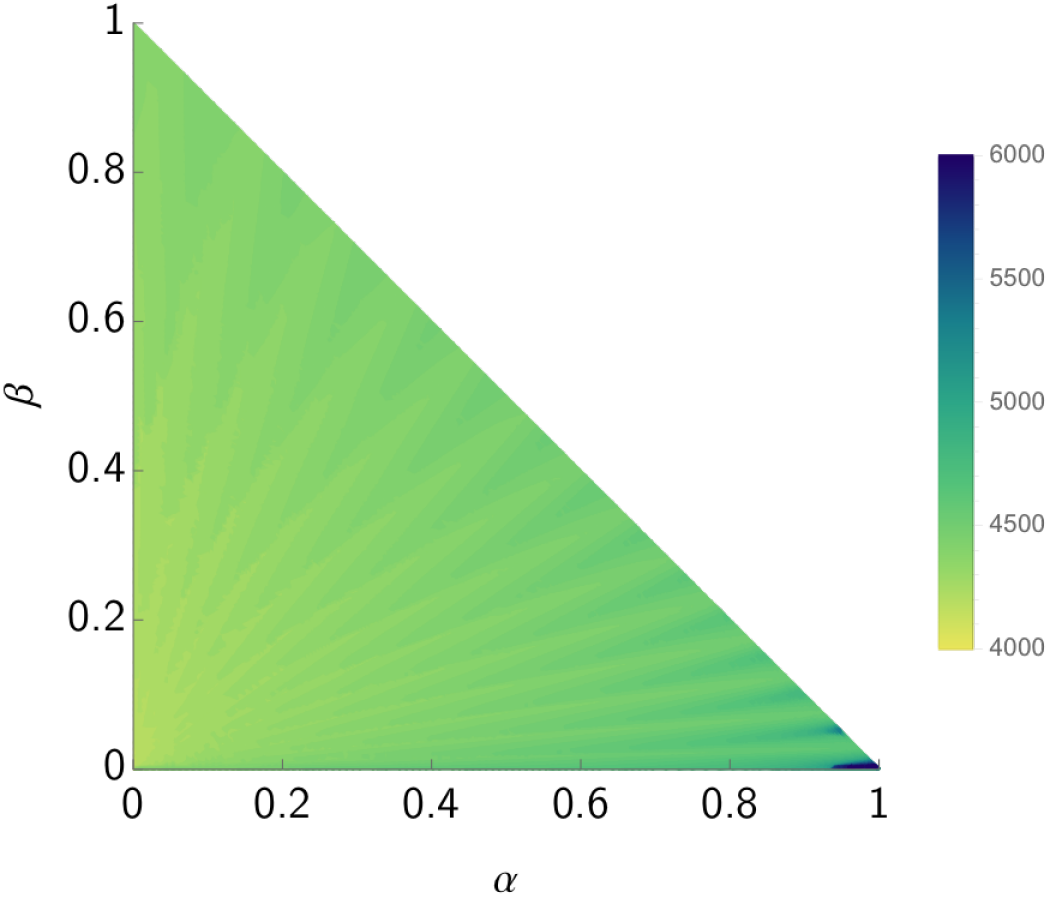
Best responders: Heat map of TCR for different values of *α* and *β*.

### 3.2. Responders

Figure 3 shows the population dynamics for the three cell types in the responder scenario. Also in this scenario, AT increases the TCR with respect to MTD for all considered assumptions on the PSA producers. Thus, the results are qualitatively similar to those of the best responders. However, quantitatively, in this scenario the TCR is much lower than in the previous case, both for MTD and AT. While applying MTD leads to a TCR of 202, the highest TCR corresponds to the cases *α* = 1, *β* = 0 and *α* = 0, *β* = 1 (499 and 497, respectively). As before, we cannot apply AT when *α* = *β* = 0. The results in terms of TCR obtained under different assumptions on *α* and *β* and TCR improvement of applying AT compared to applying MTD are displayed in Table 3.

**Figure 3:**
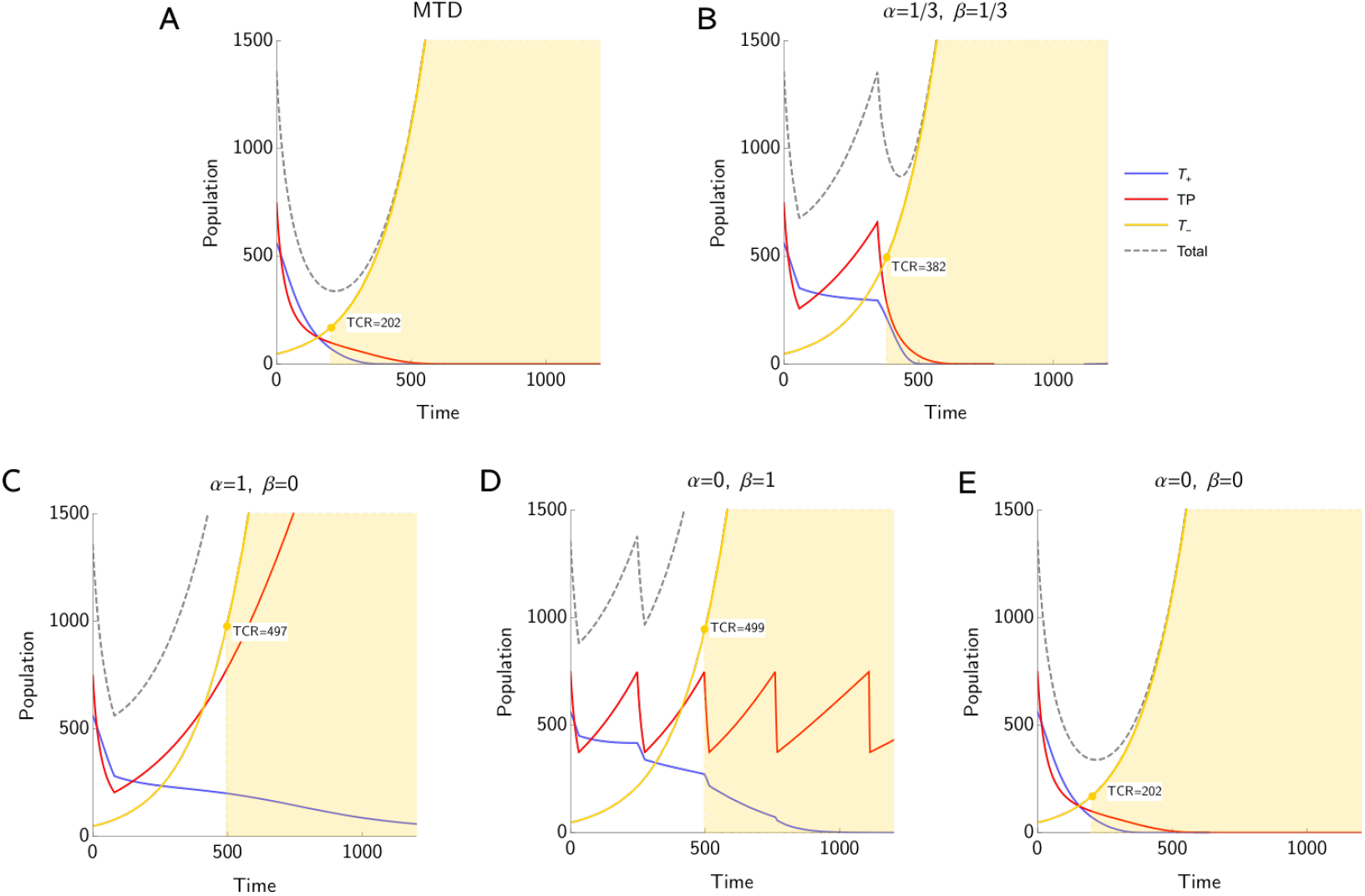
Responders: Time to competitive release under MTD (A and E) and AT for different values of *α* and *β* (B-D). For all the values of *α* and *β* considered here, AT increases the TCR compared to the standard of care with MTD. The only case where we could not apply AT is when *α* = *β* = 0, i.e., *T*^−^ cells are the only PSA producers. This is due to the fact that the treatment does not affect the PSA level in such a case. We observe the highest TCR for *α* = 0, *β* = 1, followed by the case with *α* = 1, *β* = 0, which has a similar TCR.

**Table 3:** Responders: TCR under MTD and under AT including the TCR improvement percentage depending on different assumptions on PSA production. Applying AT increases TCR in all considered cases. We observe the highest improvement in the TCR for *α* = 1, *β* = 0 and a similar improvement for *α* = 0, *β* = 1.

| Parameter values | TCR under MTD | TCR under AT | Improvement |
| --- | --- | --- | --- |
| $\alpha = 1; \beta = 0$ | 202 | 497 | 146 % |
| $\alpha = 0; \beta = 1$ | 202 | 499 | 147 % |
| $\alpha = \frac{1}{3}; \beta = \frac{1}{3}$ | 202 | 382 | 89 % |
| $\alpha = 0; \beta = 0$ | 202 | N/A | N/A |

For *α* = 1, *β* = 0, and 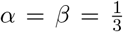, the AT treatment is stopped before reaching TCR. For *α* = 0, *β* = 1, there are at least two full treatment cycles. However, as expected, the number of cycles here is much lower than in the best responder scenario.

Figure 4 illustrates TCR for all possible values for *α* and *β*. We observe the highest values for TCR if *α >* 0.6, *β >* 0.2. Interestingly, if the *T*^+^ cells are the only PSA producers, i.e., *α* = 1, the TCR is lower. If the *T*^−^ cells contribute to the PSA production a lot, i.e., *α <* 0.2, *β <* 0.2, the TCR is the lowest.

**Figure 4:**
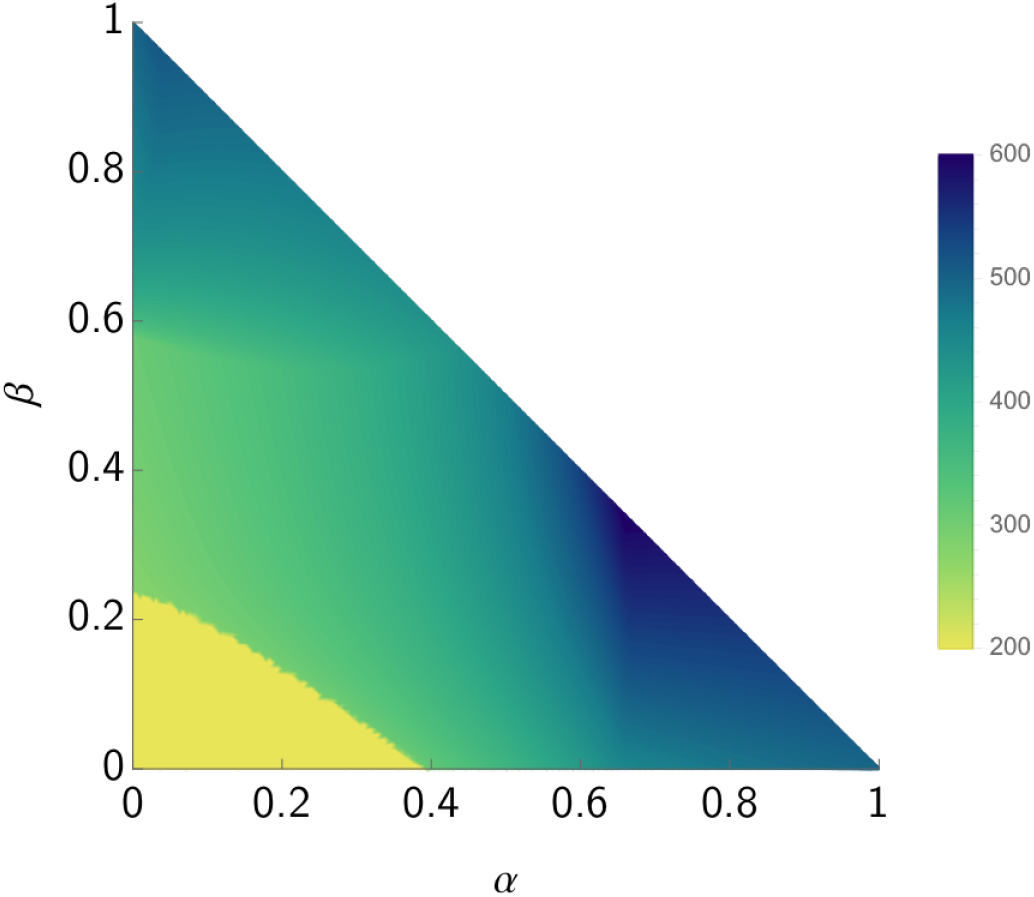
Responders: TCR heatmap with respect to different values of *α* and *β*.

### 3.3. Non-responders

Figure 5 displays the population size dynamics for *T*^*P*^, *T*^+^, and *T*^−^ cells in the non-responder scenario. As expected, the TCR is lower than the TCR of the best responder and responder scenarios. This holds under both AT and MTD. Whenever AT can be applied, i.e., whenever the treatment can be discontinued according to the treatment protocol before TCR is reached, the TCR is increased compared to the standard of care. However, for the combinations of *α* and *β* considered here, the AT protocol can be applied only if *α* = 0, *β* = 1. In this case, i.e., if *T*^*P*^ cells are the only PSA producers, AT can achieve a TCR that is about three times larger than the TCR achieved with the standard of care (see Table 4). Figure 6 supports the results displayed in Figure 5: The highest TCR can be achieved for *α* = 0, *β* = 1, while in the other three scenarios, the AT cannot be applied.

**Figure 5:**
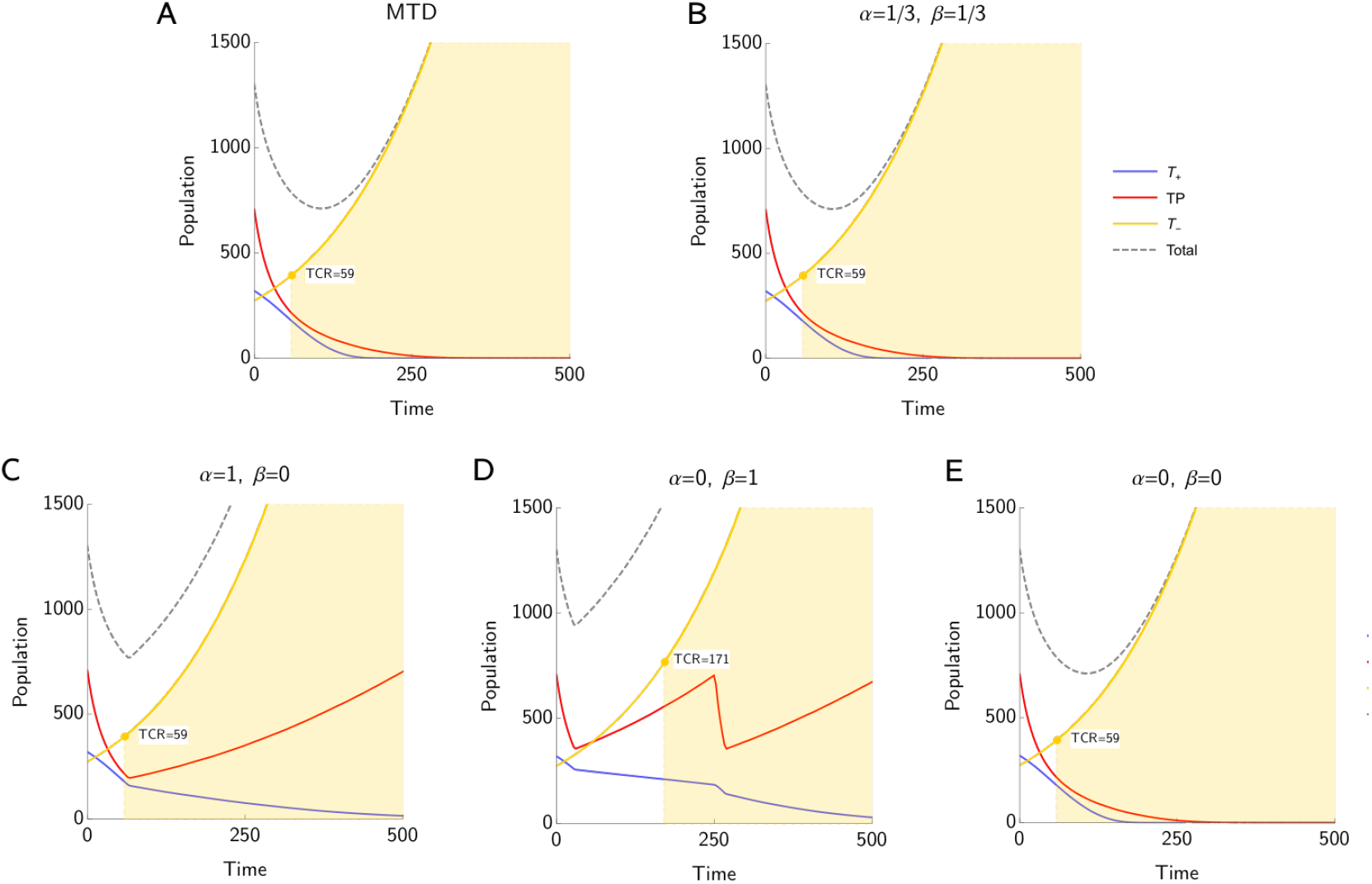
Non-responders: Time to competitive release under different assumptions on PSA production. AT can only be applied if *α* = 0, *β* = 1. In this case, we observe an increase of TCR from 59 (MTD) to 171 (AT). In all other cases, AT is not applied as the treatment cannot be discontinued before TCR.

**Table 4:** Non-responders. TCR under MTD or under AT including the TCR improvement percentage depending on different assumptions on PSA production. Only if *T*^*P*^ cells are the only cells producing PSA, AT can be applied. In this case, we observe an improvement of 188% in the TCR when applying AT instead of MTD.

| Parameter values | TCR under MTD | TCR under AT | Improvement |
| --- | --- | --- | --- |
| $\alpha = 1; \beta = 0$ | 59 | N/A | N/A |
| $\alpha = 0; \beta = 1$ | 59 | 171 | 188 % |
| $\alpha = \frac{1}{3}; \beta = \frac{1}{3}$ | 59 | N/A | N/A |
| $\alpha = 0; \beta = 0$ | 59 | N/A | N/A |

**Figure 6:**
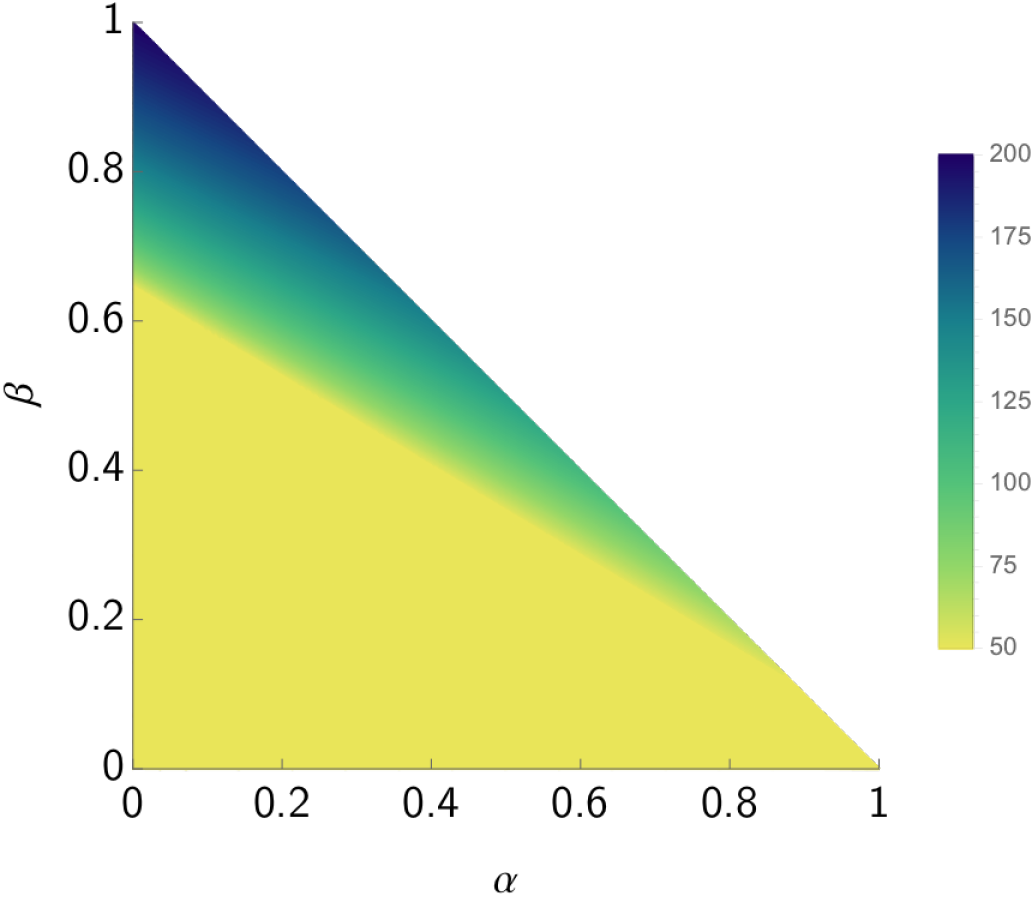
Non-responders: TCR heatmap for different values of *α* and *β*.

### 4. Discussion

PSA is a traditionally used biomarker to track the treatment-induced response in prostate cancer. It has also been used to modulate the treatment in adaptive therapy protocols, such as the one by Zhang et al. [42, 1]. However, its correlation with tumor volume is not well understood. While experimental studies revealed that the production of PSA depends on the tumor composition [18], mathematical models of adaptive therapy usually consider PSA as a surrogate for tumor burden and look only at the total cell count in order to determine when to pause or resume the treatment [42, 10, 8, 37].

Here we explored how the assumption that different cancer cell types contribute to the PSA level differently impacts the superiority of adaptive therapy protocols in metastatic castrate resistant prostate cancer (mCRPC) over the standard of care with continuous maximum tolerable dose. We focused on one particular adaptive therapy protocol, i.e. Zhang et al.’s protocol utilized in the first adaptive abiraterone therapy on mCRPC [1]. This protocol discontinues abiraterone treatment when the PSA level gets below half of its initial value and abiraterone is re-administered only once PSA recovers to its initial value [42]. We compared time to competitive release (TCR) of Zhang et al.’s adaptive protocol to that of abiraterone maximum tolerable dose when mCRPC is modelled as in [42, 10, 11]. We did this under the assumption that different cancer cell types may contribute to the PSA level differently. For example, in the limit case, we assumed that either *T*^+^ cells, *T*^*P*^ cells, or *T*^−^ cells may be the only PSA producers, while we also analyzed other scenarios, such as those when the three cell types contribute to PSA equally.

We measured time to competitive release for three categories of patients analyzed in [10]: best responders, responders and non-responders, which have no, low, and remarkable initial proportion of abiraterone-resistant *T*^−^ cells, respectively. Moreover, in the model considered here, these three categories are described by different competition matrices, corresponding to very good, intermediate and very bad prognoses, respectively.

Our results show that the Zhang et al.’s adaptive therapy protocol outperforms the standard of care based on maximum tolerable dose whenever adaptive therapy can be applied. The best responders have the longest time to competitive release (TCR) under MTD compared to the other categories. Zhang et al.’s adaptive therapy can further improve the time to competitive release by 13-32%, depending on the contribution of the different types to PSA production (Figure 1, Table 2). The greatest improvement is found in the case where the PSA is secreted almost exclusively by the *T*^+^ cells (Figure 2). When PSA is produced only by the *T*^−^ cells, Zhang et al.’s adaptive therapy cannot be applied, as demonstrated in Figure 1.

For the responders, adaptive therapy can prolong the time to competitive release by 89-147% compared to MTD (Table 3). Similarly to the previous category, the most favourable outcome corresponds to the case where the PSA is secreted (almost) only by the *T*^+^ cells, as shown in Figures 3 and 4. When *T*^−^ cells are the only type of cells contributing to the PSA production, Zhang et al.’s adaptive therapy cannot be applied.

As expected, the non-responders have the shortest time to competitive release under MTD when compared to the other categories (Table 4). Figures 5 and 6 show that for this category adaptive therapy can only be applied when the PSA is produced mostly by the *T*^*P*^ cells. In such a case, adaptive therapy can improve the TCR by 188% compared to that of the standard of care. Overall, adaptive therapy proved to lead to a higher time to competitive release than the standard of care whenever it could be applied.

Gustavsson et al. (2005) investigated PSA secretion in androgen-dependent and independent cells in vitro [18]. They reported that the level of PSA secreted by the androgen-dependent cells was tenfold higher than that by androgen-independent cells. This suggests that it might be unlikely to have the *T*^−^ cells as the main PSA producers, which corresponds to the case where adaptive therapy cannot be applied in the categories considered here. A question remains whether the coefficients *α* and *β* governing the PSA dynamics (Eq 2) can vary with time and/or tumor characteristics.

We did not consider delayed PSA dynamics, as we could not find information on how precisely the dynamics should be delayed and as we confined to the models of [10, 11] here. Identifying realistic PSA dynamics is a subject for future work.

In this work, time to competitive release measured success of the considered therapies, as opposed to a more standard time to PSA progression. While it may be difficult to estimate this time in reality, our approach is more conservative than time to PSA progression. This is because time to competitive release is typically lower than that of the PSA progression [11]. Identifying time to competitive release accurately would open a window of opportunity for patients, as physicians may have time to consider alternative treatment options to delay PSA progression once the competitive release is identified.

In this work, we considered therapy resistance as a qualitative trait, as there are two types of cancer cells (*T*^*P*^ and *T*^+^) which are targeted by the therapy and one type (*T*^−^) which does not respond, neither to androgen deprivation nor abiraterone treatment as it is independent of testosterone. Conversely, some recent works considered quantitative resistance [31, 27, 38]. In any case, even with resistance as a quantitative trait, Zhang’s treatment protocol would be more successful than MTD [26, 28].

Since Zhang et al.’s trial, different evolutionary cancer treatment protocols, i.e. protocols that anticipate and steer eco-evolutionary cancer dynamics, have been proposed in the literature [36, 27, 26, 15, 38, 31, 37]. For example, Viossat and Noble (2021) demonstrated that already discontinuing the treatment when PSA reduces to a higher proportion (for example 80%) of its initial value would be more effective than the original Zhang et al.’s protocol where treatment is continued until PSA drops by half. Cunningham et al (2020) demonstrated that the mCRPC tumor burden may be stabilized through a dose titration protocol [11]. Gatenby et al (2019) suggested that cure in mCRPC may be possible if a different therapy is applied in a strategic way when the initial treatment response is observed [16]. The fact that the classical adaptive therapy protocol that we analyzed here performs very well under the vast majority of assumptions on the contribution od different cell types’ to the prostate specific antigen level is a very good news for both these other evolutionary therapies and patients with metastatic disease, as long as PSA remains the main marker for tumor progression. To further improve therapy design in mCRPC, a better understanding of mechanisms behind the PSA production and/or alternative biomarkers in mCRPC are needed. Combining these with our predictive mathematical models will help to answer key questions in the ecology and evolution of cancer, such as “How can game theory be utilized to understand tumorigenesis and potentially guide therapy?” or “Are there measures of the evolution and ecology of tumours that can be used to develop a classification system for tumours, so as to improve prediction, prognosis and management of tumours?” [13]. More importantly, such a development will likely improve odds for patients suffering the metastatic disease.

## Acknowledgements

We thank Dr. Heiko Enderling and Dr. Renee Brady-Nicholls for their valuable feedback and for pointing out interesting related work. This research was supported by European Union’s Horizon 2020 research and innovation programme under the Marie Skłodowska-Curie grant agreement number 955708 and the Dutch National Foundation projects ENWPR.020.006 and OCENW.KLEIN.277.

## References

[1] H. Lee Moffitt Cancer Center and Research Institute: Adaptive abiraterone therapy for metastatic castration resistant prostate cancer (NCT02415621).

[2] S. P. Balk, Y.-J. Ko, and G. J. Bubley. Biology of prostate-specific antigen. Journal of clinical oncology, 21(2):383–391, 2003.

[3] J. P. Barnaby, I. C. Sorribes, and H. V. Jain. Relating prostate-specific antigen leakage with vascular tumor growth in a mathematical model of prostate cancer response to androgen deprivation. Computational and Systems Oncology, 1(2):e1014, 2021.

[4] D. Berthold, G. Pond, R. de Wit, M. Eisenberger, I. F. Tannock, et al. Survival and psa response of patients in the tax 327 study who crossed over to receive docetaxel after mitoxantrone or vice versa. Annals of oncology, 19(10):1749–1753, 2008.

[5] R. Brady-Nicholls, J. D. Nagy, T. A. Gerke, T. Zhang, A. Z. Wang, J. Zhang, R. A. Gatenby, and H. Enderling. Prostate-specific antigen dynamics predict individual responses to intermittent androgen deprivation. Nature communications, 11(1):1–13, 2020.

[6] R. Brady-Nicholls, J. Zhang, T. Zhang, A. Z. Wang, R. Butler, R. A. Gatenby, and H. Enderling. Predicting patient-specific response to adaptive therapy in metastatic castration-resistant prostate cancer using prostate-specific antigen dynamics. Neoplasia, 23(9):851–858, 2021.

[7] N. Bruchovsky, P. S. Rennie, A. J. Coldman, S. L. Goldenberg, M. To, and D. Lawson. Effects of androgen withdrawal on the stem cell composition of the shionogi carcinoma. Cancer research, 50(8):2275–2282, 1990.

[8] J. Cunningham. A call for integrated metastatic management. Nature Ecology & Evolution, 2019.

[9] J. J. Cunningham. Evolutionary Game Theory and Optimal Control for Integrated Metastatic Management of Prostate Cancer. PhD thesis, Maastricht University, Maastricht, The Netherlands, September 2021.

[10] J. J. Cunningham, J. S. Brown, R. A. Gatenby, and K. Staňková. Optimal control to develop therapeutic strategies for metastatic castrate resistant prostate cancer. Journal of theoretical biology, 459:67–78, 2018.

[11] J. J. Cunningham, F. Thuijsman, R. Peeters, Y. Viossat, J. S. Brown, R. A. Gatenby, and K. Staňková. Optimal control to reach eco-evolutionary stability in metastatic castrate resistant prostate cancer. PLOS ONE, 15(12):1–24, 2020.

[12] J. S. De Bono, C. J. Logothetis, A. Molina, K. Fizazi, S. North, L. Chu, K. N. Chi, R. J. Jones, O. B. Goodman Jr, F. Saad, et al. Abiraterone and increased survival in metastatic prostate cancer. New England Journal of Medicine, 364(21):1995–2005, 2011.

[13] A. M. Dujon, A. Aktipis, C. Alix-Panabières, S. R. Amend, A. M. Boddy, J. S. Brown, J.-P. Capp, J. DeGregori, P. Ewald, R. Gatenby, M. Gerlinger, M. Giraudeau, R. K. Hamede, E. Hansen, I. Kareva, C. C. Maley, A. Marusyk, N. McGranahan, M. J. Metzger, A. M. Nedelcu, R. Noble, L. Nunney, K. J. Pienta, K. Polyak, P. Pujol, A. F. Read, B. Roche, S. Sebens, E. Solary, K. Staňková, H. Swain Ewald, F. Thomas, and B. Ujvari. Identifying key questions in the ecology and evolution of cancer. Evolutionary Applications, 0000:1–16, 2020. https://note.org/10.1111/eva.13190.

[14] R. Gatenby, A. Silva, R. Gillies, and B. Frieden. Adaptive therapy. Cancer Research, 69(11):4894–4903, 2009.

[15] R. A. Gatenby, J. Zhang, and J. S. Brown. First strike–second strike strategies in metastatic cancer: Lessons from the evolutionary dynamics of extinction. Cancer Research, 79(13):3174–3177, 2019.

[16] R. A. Gatenby, J. Zhang, and J. S. Brown. First strike–second strike strategies in metastatic cancer: Lessons from the evolutionary dynamics of extinction. Cancer Research, 2019.

[17] Q. Guo, Z. Lu, Y. Hirata, and K. Aihara. Parameter estimation and optimal scheduling algorithm for a mathematical model of intermittent androgen suppression therapy for prostate cancer. Chaos: An Interdisciplinary Journal of Nonlinear Science, 23(4):043125, 2013.

[18] H. Gustavsson, K. Welén, and J.-E. Damber. Transition of an androgen-dependent human prostate cancer cell line into an androgen-independent subline is associated with increased angiogenesis. The Prostate, 62(4):364–373, 2005.

[19] E. Hansen and A. F. Read. Modifying adaptive therapy to enhance competitive suppression. Cancers, 12(12):3556, 2020.

[20] Y. Hirata, N. Bruchovsky, and K. Aihara. Development of a mathematical model that predicts the outcome of hormone therapy for prostate cancer. Journal of theoretical biology, 264(2):517–527, 2010.

[21] A. M. Ideta, G. Tanaka, T. Takeuchi, and K. Aihara. A mathematical model of intermittent androgen suppression for prostate cancer. Journal of nonlinear science, 18(6):593, 2008.

[22] R. B. Kato, V. Srougi, F. A. Salvadori, P. P. M. R. Ayres, K. M. Leite, and M. Srougi. Pretreatment tumor volume estimation based on total serum psa in patients with localized prostate cancer. Clinics, 63(6):759–762, 2008.

[23] J. W. Moul, R. R. Connelly, R. M. Mooneyhan, W. Zhang, I. A. Sesterhenn, F. Mostofi, and D. G. McLeod. Racial differences in tumor volume and prostate specific antigen among radical prostatectomy patients. The Journal of urology, 162(2):394–397, 1999.

[24] J. E. Oesterling, W. H. Cooner, S. J. Jacobsen, H. A. Guess, and M. M. Lieber. Influence of patient age on the serum psa concentration. an important clinical observation. The Urologic clinics of North America, 20(4):671–680, 1993.

[25] C. Pezaro, H. H. Woo, and I. D. Davis. Prostate cancer: measuring psa. Internal medicine journal, 44(5):433–440, 2014.

[26] M. Pressley, M. Salvioli, D. B. Lewis, C. L. Richards, J. S. Brown, and K. Staňková. Evolutionary dynamics of treatment-induced resistance in cancer informs understanding of rapid evolution in natural systems. Frontiers in Ecology and Evolution, 9:460, 2021.

[27] D. R. Reed, J. Metts, M. Pressley, B. L. Fridley, M. Hayashi, M. S. Isakoff, D. M. Loeb, R. Makanji, R. D. Roberts, M. Trucco, et al. An evolutionary framework for treating pediatric sarcomas. Cancer, 126(11):2577–2587, 2020.

[28] M. Salvioli, H. Garjani, J. S. Brown, J. Dubbeldam, and K. Staňková. Stackelberg evolutionary games of cancer treatment: Tumor stabilization as an alternative to dynamic treatment protocols. submitted, 2020.

[29] A. J. Schrader, M. Boegemann, C.-H. Ohlmann, T. J. Schnoeller, L.-M. Krabbe, T. Hajili, F. Jentzmik, M. Stoeckle, M. Schrader, E. Herrmann, et al. Enzalutamide in castration-resistant prostate cancer patients progressing after docetaxel and abiraterone. European urology, (1):30–36, 2014.

[30] T. Shimada and K. Aihara. A nonlinear model with competition between prostate tumor cells and its application to intermittent androgen suppression therapy of prostate cancer. Mathematical biosciences, 214(1-2):134–139, 2008.

[31] K. Staňková, J. S. Brown, W. S. Dalton, and R. A. Gatenby. Optimizing cancer treatment using game theory: A review. JAMA Oncology, 5(1):96–103, 2019.

[32] T. Suzuki, N. Bruchovsky, and K. Aihara. Piecewise affine systems modelling for optimizing hormone therapy of prostate cancer. Philosophical Transactions of the Royal Society A: Mathematical, Physical and Engineering Sciences, 368(1930):5045–5059, 2010.

[33] K. R. Swanson, L. D. True, D. W. Lin, K. R. Buhler, R. Vessella, and J. D. Murray. A quantitative model for the dynamics of serum prostate-specific antigen as a marker for cancerous growth: an explanation for a medical anomaly. The American journal of pathology, 158(6):2195–2199, 2001.

[34] G. Tanaka, Y. Hirata, S. L. Goldenberg, N. Bruchovsky, and K. Aihara. Mathematical modelling of prostate cancer growth and its application to hormone therapy. Philosophical Transactions of the Royal Society A: Mathematical, Physical and Engineering Sciences, 368(1930):5029–5044, 2010.

[35] Y. Tao, Q. Guo, and K. Aihara. A partial differential equation model and its reduction to an ordinary differential equation model for prostate tumor growth under intermittent hormone therapy. Journal of mathematical biology, 69(4):817–838, 2014.

[36] Y. Viossat and R. Noble. A theoretical analysis of tumour containment. Nature Ecology and Evolution, 2021.

[37] J. West, M. Dinh, J. Brown, J. Zhang, A. Anderson, and R. Gatenby. Multidrug cancer therapy in metastatic castrate-resistant prostate cancer: An evolution-based strategy. Clinical Cancer Research, 25(14):4413–4421, 2019.

[38] B. Wölfl, H. te Rietmole, M. Salvioli, A. Kaznatchev, F. Thuijsman, J. S. Brown, B. Burgering, and K. Staňková. The contribution of evolutionary game theory to understanding and treating cancer. medRxiv, 2020.

[39] L. You, J. S. Brown, F. Thuijsman, J. J. Cunningham, R. A. Gatenby, J. Zhang, and K. Staňková. Spatial vs. non-spatial eco-evolutionary dynamics in a tumor growth model. Journal of Theoretical Biology, 435:78–97, 2017. 10.1016/j.jtbi.2017.08.022.

[40] J. Zhang, J. Cunningham, J. Brown, and R. Gatenby. Integrating evolutionary dynamics into treatment of metastatic castrate-resistant prostate cancer. Nat. Commun., 8, 2017.

[41] J. Zhang, J. J. Cunningham, J. S. Brown, , and R. A. Gatenby. Evolutionary dynamics of combined LHRH analogs and ar directed agents in first line therapy of metastatic castrate sensitive prostate cancer. submitted.

[42] J. Zhang, J. J. Cunningham, J. S. Brown, and R. A. Gatenby. Integrating evolutionary dynamics into treatment of metastatic castrate-resistant prostate cancer. Nature communications, 8(1):1816, 2017.

